# If you give a mouse a poopsicle: a novel fecal microbiota transplant method for exploring the role of the gut microbiome in stress-related outcomes in mice

**DOI:** 10.64898/2026.02.16.705192

**Authors:** Monica A. Tschang, Ronin Deo-Campo Vuong, Baylee Eilers, Denise Chac, Adam Waalkes, Kelsi Penewit, Alyssa Easton, Bryan Schuessler, Renata Daniels, Ana A. Weil, Stephen J. Salipante, Sean M. Gibbons, Abigail G. Schindler

## Abstract

The microbiome-gut-brain axis is a mediator of stress-related disorders. The number of preclinical studies exploring the potential causal mechanism of this connection using fecal microbiota transplantation (FMT) is growing. However, the most common method for delivering fecal transplants in rodent models is still oral gavage, which creates an adverse experience that may confound stress-related outcomes. Here, we establish an alternative methodology for FMT that decreases stress induced by traditional experimental procedures. We first used preference and anxiety behavior assays to identify antibiotic therapies having maximal tolerability and minimal anxiolytic properties. We then collected feces from donor mice and homogenized them with a microbe-stabilizing buffer to create a slurry, which was frozen into pellets (“poopsicles”) for subsequent FMT. Recipient mice voluntarily consumed the pellets, and blood was collected to compare corticosterone levels relative to traditional gavage FMT. Plasma corticosterone levels were found to be significantly lower in mice receiving FMT via pellets compared to oral gavage. Furthermore, relative to gavage FMT, microbial signatures of mice receiving FMT via pellets were more similar to those of the donor pellets at one week following final FMT and were sustained for up to six weeks, as assessed by comparing Bray-Curtis beta-diversity distances. Together, these results establish effective antibiotic and FMT methods that minimize treatment-induced stress, while effectively transplanting fecal microbes between murine conspecifics.

## 1 Introduction

The gut microbiome is an increasingly popular target for treating and understanding various health conditions, including stress (Bashir and Khan, 2022; Loh *et al*., 2024; Margolis, Cryan and Mayer, 2021; Wiley *et al*., 2017). Indeed, multiple studies report associations between gut microbiome composition and psychiatric disorders, including depression, social anxiety disorder, and post-traumatic stress disorder (Bersani *et al*., 2020; Butler *et al*., 2023; Cruz-Pereira *et al*., 2020; Tillmann *et al*., 2019). Through multiple bidirectional avenues of communication between the gastrointestinal tract and the brain, gut-residing microbes can influence the central nervous system (Margolis, Cryan and Mayer, 2021). As further evidence emerges to support the connection between behavioral neuroscience and microbial ecology, our understanding of the biological basis of neuropsychiatric conditions is greatly enriched.

Despite its potential, the bidirectionality of the microbiota-gut-brain axis presents a challenging environment where it is difficult to distinguish cause from effect (Margolis, Cryan and Mayer, 2021; Rengasamy *et al*., 2021). Thus, a significant hurdle in clinically-focused microbiome research is establishing causality, because most human microbiome studies are merely correlational (Lynch, Parke and O’Malley, 2019; Walter *et al*., 2020). In rodent preclinical models, the favored strategy to address this hurdle is to use a fecal microbiota transplant (FMT), which transfers the microbiome of one subject to another, isolating the effect of the gut microbiome on health outcomes (Gheorghe *et al*., 2021; Keubler *et al*., 2023). A cornerstone study that established FMT in the gut-brain-axis field was performed by transferring feces from patients with depression to microbe-depleted rats, which induced characteristics of depression in the recipient animals (Kelly *et al*., 2016).

While FMT is not a new approach (D and Venkatesh, 2023), there remains a lack of standardization across multiple experimental domains required for successful FMT in murine studies (Mingaila, Atzeni and Burokas, 2023; Secombe *et al*., 2021) (Table 1). Systematic reviews highlight that inconsistent protocols, including differences in donor selection, recipient preparation (e.g., antibiotic vs. germ-free), administration route, dosing interval, and handling conditions, lead to variable engraftment and behavioral outcomes, impeding robust interpretation and cross-study comparison (Bokoliya *et al*., 2021; Keubler *et al*., 2023; Mingaila, Atzeni and Burokas, 2023). For example, the first step of the FMT is to find or create an appropriate host suitable for accommodating the transplanted microbiome. This usually involves either using gnotobiotic (germ-free) mice or broad-spectrum antibiotics to ensure that the native microbiota does not block engraftment of the transplanted microbiota. Antibiotics are the more accessible option for laboratories without high-cost germ-free facilities; however, the administration of antibiotics varies in dosage, length, type, and route of administration across studies (Bokoliya *et al*., 2021; Kennedy, King and Baldridge, 2018).

**Table 1:** Dosing schemes for oral gavage FMT from rodent conspecifics in stress-related behavioral models.

| Disorder Investigated | Animal Model | Timeline/Duration | Microbial Content | Dose | Reference |
| --- | --- | --- | --- | --- | --- |
| Chronic Stress | mouse | 1 and 4 days after antibiotic discontinuation<br>(2 doses total) | 1 mg feces in 5 mL PBS (phosphate buffer saline) | 300 uL | (Siopi *et al.*, 2020) |
| Depression | mouse | 1x/day for 3 consecutive days<br>(3 doses total) | ~2 × 10<sup>8</sup> viable probiotic fecal bacteria in PBS | 300 uL | (Zhang *et al.*, 2019) |
| Chronic unpredictable mild stress | mouse | 1x/2 days for 14 days (7 doses) | Supernatant of cecal content diluted 40-fold in PBS | 200 uL | (Li *et al.*, 2019) |
| Alcohol withdrawal-induced anxiety | mouse | 1x/day for 14 consecutive days<br>(14 doses total) | Supernatant of 0.1g feces in 1mL saline | 200 uL | (Xiao *et al.*, 2018) |
| Depression | rat | 1x/3 days for 16 days<br>(6 doses total) | 1 g feces in 5 mL saline | 750 uL | (Tillmann *et al.*, 2019) |
| Depression | rat | 1x for 3 days (3 doses) | 1g feces in 15 mL PBS | 200 uL | (Lv *et al.*, 2019) |
| Social defeat | rat | 1x/5 days (5 doses) | 5 mg feces in 1 mL PBS | 2 mL | (Pearson-Leary *et al.*, 2020) |

Most concerningly, many studies reported poor consumption of antibiotics administered ad libitum, likely due to the especially bitter or otherwise unpalatable taste of many antibiotics (Almonte *et al*., 2022; Hill *et al*., 2010; Rath *et al*., 2001; Reikvam *et al*., 2011; Zákostelská *et al*., 2016). Dehydration as a side-effect of mice disliking the antibiotics dissolved in their water poses a physiological concern as well as risk of altering gut microbiota composition (Sato *et al*., 2024). Dosing antibiotics via oral gavage, which usually requires two doses a day for consistent microbiota depletion, circumvents palatability issues and provides more experimental control (Hill *et al*., 2010; Reikvam *et al*., 2011; Tirelle *et al*., 2020). Oral gavage is also the primary method of delivery of FMT in rodent models (Table 1).

However, the gavage process itself significantly elevates plasma corticosterone levels 30 minutes after dosing (Gonzales *et al*., 2014) and studies suggest that this elevation could underlie the overall lack of behavioral changes in studies following FMT in rodents (Tillmann et al., 2019). Consequently, for studies with stress as a behavioral outcome, data may be further confounded by the mode of FMT delivery in addition to the lack of standardization in FMT preparation. Given that mice evolved to consume their own and conspecifics’ feces in a behavior known as coprophagy, which important in maintaining rodent health and immunity (Sha *et al*., 2023), voluntary consumption of antibiotics or fecal content presents as a less stress-inducing, more desirable option for FMT.

Several alternative strategies to oral gavage that leverage voluntary intake via peanut butter (Gonzales *et al*., 2014), strawberry jam (Teixeira-Santos, Albino-Teixeira and Pinho, 2021), chocolate hazelnut spread (Abelson *et al*., 2012; Jacobsen *et al*., 2011; Kalliokoski *et al*., 2011), bacon-flavored transgenic dough (Walker *et al*., 2012), and gelatin (Hovard *et al*., 2015; Abraham *et al*., 2020). However, all of these alternative methods were used to deliver oral drugs, and not biological substrates or live cultures, as is the case for FMTs. In sum, better preclinical models of FMT in mice are needed to improve the experimental validity of gut-brain axis research, particularly in stress-related disorders. Therefore, we aimed to optimize and validate two experimental components of the FMT procedure for researchers using murine models to study stress: 1) an antibiotic regimen that effectively eliminates native gut flora without inducing aversion or anxiety-like behaviors, and 2) an FMT delivery method that successfully transfers microbiota characteristics from one mouse to another and relies on self-administration rather than restraint and gavage.

## 2 Materials and Methods

### 2.1 Animals

All experiments utilized male C57Bl/6J mice aged 9–11 weeks old at the time of arrival from Jax (Sacramento, CA) to Veterans Affairs (VA) Puget Sound Health Care System (Seattle, WA). Mice were co-housed three to four per cage on a 12:12 light:dark cycle (lights on at 06:00) and were given ad libitum food and water unless otherwise indicated. Room temperature was 21°C; room humidity was 35%. All animal experiments were carried out in accordance with Association for Assessment and Accreditation of Laboratory Animal Care guidelines and were approved by the VA Puget Sound Institutional Animal Care and Use Committees. Mice were acclimated to the housing room for a week following arrival and subsequently handled for an additional week prior to experimentation.

### 2.2 Antibiotics

VNAM: 0.5 mg/mL vancomycin (Sigma Aldrich) 1 mg/mL neomycin (Sigma Aldrich), 1 mg/mL ampicillin (Sigma Aldrich), and 1mg/mL of metronidazole (Sigma Aldrich) were dissolved in 1 L of sterilized water from the Animal Research Facility. Amphotericin-β (Sigma Aldrich) was sonicated in 10mL sterilized water for five minutes to encourage solubilization before combining with the rest of the solution for a total of 0.01□mg/mL amphotericin-β (Sigma Aldrich). After 30 minutes of stirring at room temperature, the antibiotic solution was aliquoted into 50 mL Falcon tubes before freezing at −80 °C. Before use, aliquots were thawed and the tube caps were replaced with sipper tops. Antibiotics were administered ad libitum to each home cage, changing bottles every two days.

VNAA: Identical antibiotics included in VNAM except without the metronidazole

VNBP: 0.5 mg/mL vancomycin, 2 mg/mL neomycin, 0.5 mg/L bacitracin, and 1.2 μg/mL pimaricin were dissolved in 1 L of sterilized water for 30 minutes at room temperature. Subsequently, the antibiotic solution was distributed into 50 mL aliquots before freezing at −80°C. Aliquots were thawed before use.

In the two-bottle choice experiment, 50 mL aliquots of both antibiotics (VNAA and VNBP) were given to each cage ad libitum, changing bottles every 2 days. Mice and antibiotic bottle weights were measured daily throughout the study to monitor body weight retention and liquid consumption.

### 2.3 Open Field Test

Mice were tested for locomotor deficits and anxiety-like behaviors in an open field assay prior to, 1 day, and 4 days post antibiotic treatment. Mice were placed in a large (1 m in diameter), open, circular arena split into center and outer sections for 5 minutes. Speed, total distance traveled, time spent in the center of the field, time delay to first entry to the center, and the number of entries into the center, were recorded from above and analyzed with Anymaze (Wood Dale, IL). Less time spent in the center of the arena was indicative of an anxiety-like phenotype.

### 2.4 Fecal Collection

Mice were taken out of their home cages and temporarily single-housed in an empty sterile cage until defecation. Fresh feces were collected from mice between 08:00-11:00 am using sterile technique. Feces were flash-frozen in liquid nitrogen and stored at −80 °C until shipment to Diversigen or a separate facility at the University of Washington for downstream processing and analysis.

### 2.5 Fecal Microbiome Analysis

Shallow shotgun sequencing was conducted by Diversigen Inc. (New Brighton, MN). DNA was extracted from feces and sequenced using the provider’s BoosterShot Shallow Shotgun Sequencing protocol. DNA sequences were aligned to a curated database containing all representative bacterial genomes in RefSeq. Alignments were made at 97% identity against all reference genomes. Every input sequence was compared to every reference sequence in the Diversigen Venti database using fully gapped alignment with BURST (Al-Ghalith and Knights, 2020). To assign taxonomy, each input sequence was assigned the lowest common ancestor that was consistent across at least 80% of all reference sequences tied for best hit. The number of counts for each operational taxonomic unit (OTU) was normalized to the OTU’s genome length. Samples with <10,000 sequences were discarded. OTUs accounting for less than one millionth of all strain-level markers and those with <0.01% of their unique genome regions covered (and < 0.1% of the whole genome) at the species level were discarded.

16S rRNA sequencing was conducted by the Microbial Interactions & Microbiome Center (mim_c) at the University of Washington. DNA was extracted from fecal samples using QIAamp PowerFecal Pro DNA Kit (Qiagen, Hilden, Germany). Amplicon libraries of the 16S rRNA gene V3-V4 region were prepared using primers 347F and 803R (Klindworth *et al*., 2013) in accordance with Illumina’s recommendations for 16S Metagenomic Sequencing Library Preparation (Part# 15044223 Rev. B). Libraries were sequenced using Illumina NextSeq2000® with 600-cycle X-LEAP chemistries. After demultiplexing, paired-end sequences were imported into QIIME2 (v. 2023.9.1) (Bolyen *et al*., 2019). Primers were trimmed using the cutadapt plugin (v 4.5) (Kechin *et al*., 2017). Denoising, quality filtering, and enumeration of amplicon sequence variants (ASVs) were performed using DADA2 (Callahan *et al*., 2016). Taxonomic assignments were established using scikit-learn naïve Bayes classifier trained on the SILVA 128 release reference tree, which was also used to construct a SATé-Enabled Phylogenetic Placement (SEPP) phylogenetic tree of the sequences (Janssen *et al*., 2018) for downstream analysis. Beta diversity was calculated in QIIME2 using the core phylogenetics metrics plugin.

### 2.6 Pellet (Poopsicle) Preparation

Feces were collected from donor mice likewise using sterile technique. However, instead of flash-freezing after collection, feces were combined and homogenized with chilled and filtered Brain Heart Infusion (BHI) media with 30% glycerol and chilled DietGel Recovery at a 1:9:4 ratio by weight using homogenization beads and a Bead Genie (Scientific Industries Inc, Bohemia, NY) at room temperature. This fecal homogenate was distributed into 200 μL aliquots that were then stored at −80°C until use. Each frozen pellet, also called a poopsicle, contains ∼0.015 g feces.

### 2.7 Pellet Habituation and Fecal Microbiota Transplant (FMT)

Frozen homogenate control (“blank”) pellets were created by following the preparation above, but excluded feces. Additional DietGel Recovery was added to make up for the fecal matter weight for a final ratio of 9:5 glycerol-BHI media to DietGel Recovery. Recipient mice were habituated to consuming blank pellets a week prior to the FMT. During habituation, mice were individually put in empty, sterile cages along with two frozen blank pellets were placed in the cages for mice to consume voluntarily for 30 minutes. Control pellets were put on sterile weigh boats and weighed before and after the eating period. On average, mice ate 0.164 ± 0.013 g of the blank pellets. For the subsequent FMT, antibiotics (VNAA) were given for 4 days in home cages ad libitum followed by a washout day in which mice drank water before pellet exposure. FMT pellets were likewise put on plastic weigh boats for pre- and post-weight measurements and for ease of delivery into cages. On average, mice ate 0.182 ± 0.009 g of the FMT pellets. Mice were given free access to the pellets for two hours or until they finished them, after which they were returned to their home cages.

### 2.8 High-Fat/High-Sugar Diet for Donors

Assigned donor mice (n= 16, co-housed in cages of four) received a high-fat, high-sugar dietary supplement in the form of a cup of DietGel Boost in their home cages in addition to their standard water and chow. The DietGel Boost cup was monitored daily for 6 days, after which we collected and pooled feces from all donor mice to be made into pellets for FMT as described above.

### 2.9 Oral Gavage

Mice were temporarily restrained to administer 200 mL of a control slurry (same composition as the blank pellet but not frozen) to each mouse via a 22-gauge, 1.5-inch metal gavage needle between 9:00 and 11:00 am for three consecutive days.

### 2.10 Corticosterone

30 minutes following final FMT or antibiotic administration, mice were euthanized via cervical decapitation. Trunk blood was collected in serum-separator tubes, allowed to clot for 30-50 minutes, and then centrifuged at 3,000 x g for 10 minutes at room temperature. Afterwards, serum was aliquoted and stored at −80 °C until analysis. Corticosterone levels were then analyzed using a Corticosterone Multi-Format ELISA Kit (Arbor Assays, Ann Arbor, MI). Blood samples were all collected before noon.

### 2.11 Anaerobic Plating

Fecal, pellet, and DietGel samples were introduced into anaerobic environments and thawed. Fecal samples were split in half and placed into a new 1.5 mL tube. The remaining samples were vortexed. 50 μL of samples were plated as a lawn onto TSA + 5% sheep’s blood (TSAb) and incubated at 37°C for 72 hours. Additional samples were plated at lower dilutions (either 1:10 or 1 loop of bacteria), depending on the sample’s viscosity. Bacterial growth on TSAb plates was scraped into a 1.5 mL tube and centrifuged to pellet bacteria for sequencing.

### 2.12 Statistical Analysis

Data were expressed as mean ± SEM. Lines of best fit were calculated with simple linear regression. Differences between groups were determined using a two-tailed Student’s t-test, one-way analysis of variance (ANOVA), or two-way (repeated measures when appropriate) analysis of variance (ANOVA) followed by post hoc testing using either Šídák or Tukey’s multiple comparisons test. Bray-Curtis dissimilarity was used to estimate beta diversity among samples and represented using PCoA plots. Permutational Multivariate Analysis of Variance (PERMANOVA) was used to compare beta diversity across groups (999 permutations). Reported p-values denote two-tailed probabilities of p ≤ 0.05, and non-significance (ns) indicates p > 0.05. Statistical analysis and visualization were conducted using GraphPad Prism 9.0 (GraphPad Software, Inc., La Jolla, CA) and with custom R scripts.

## 3 Results

### 3.1 Mice maintain body weight during VNAA exposure and do not differ from water controls

One of the most common antibiotic cocktails to deplete native gut microbiota prior to FMT is comprised of vancomycin, neomycin, ampicillin, amphotericin-β, and metronidazole (VNAM) (Figure 1Ai). However, administration of VNAM ad libitum in the drinking water resulted in over 20% weight loss in less than a week, due to severe aversion (n = 21; Simple linear regression: b=-0.045, R2=.879, F[1,103]=751, p<0.0001) (Figure 1B - blue circles). We therefore compared the same combination minus the metronidazole (VNAA) to an alternative antibiotic combination containing vancomycin, neomycin, bacitracin, and pimaricin (VNBP) (Hofford *et al*., 2021; Kiraly *et al*., 2016) in a two-bottle choice test performed in the home cage for two weeks (Figure 1Aii). Mice in these cages gained body weight (n = 9; Simple linear regression: b = 0.004, R2=.274, F[1,124]=46.7, p<0.0001) (Figure 1B - purple squares). Comparing the first six days between VNAM and VNAA/VNBP, mice exposed to VNAM lost more weight compared to mice exposed to VNAA and VNBP (2-way RM ANOVA: interaction effect F[4,112]=196, p<0.0001, main effect of time F[2, 58]=187, p<0.0001, and main effect of antibiotic solution F[1, 28]=197, p<0.0001). Šídák’s multiple comparison post-hoc test revealed that the percent body weight of VNAM-exposed mice was significantly lower than that of VNAA/VNBP-exposed mice starting at day two until day six (p<0.0001) (Figure 1B) (note VNAM was discontinued after day six due to severe weight loss due to dehydration). Mice preferred VNAA to VNBP in the two-bottle choice test across the two-week exposure when comparing the average daily amount consumed of each solution by each cage, normalized to the combined body weight in the cage (Student’s paired t test: t[2]=4.63, p=0.044) (Figure 1C).

**Figure 1.**
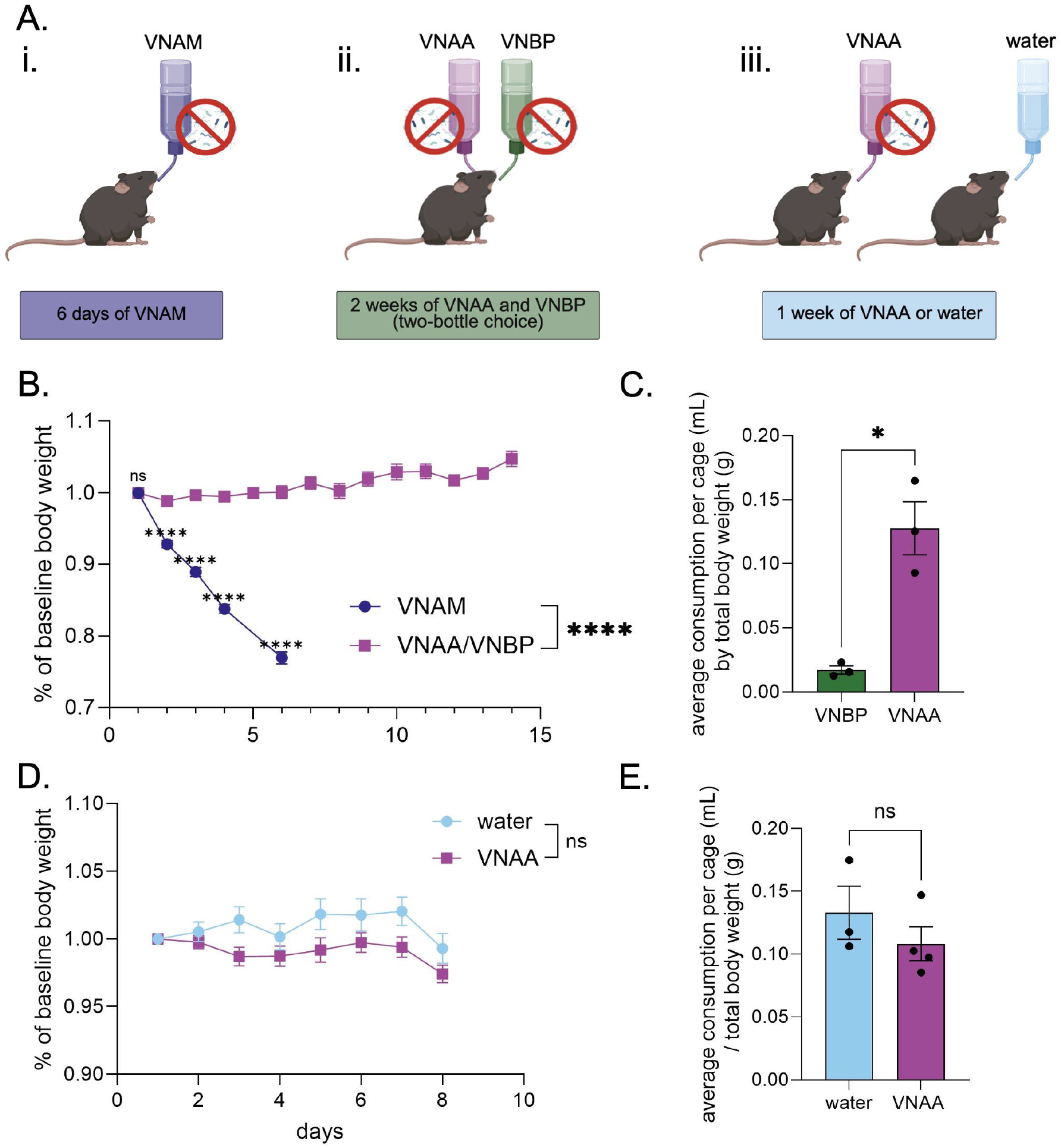
An alternative antibiotic cocktail does not reduce body weight. A: Experimental design of three different antibiotic regimens. B and D: timeline of bodyweight as a percentage of the starting body weight over the course of VNAM, or VNAA/VNBP (b) and water or VNAA (d). C and E: Average volume of administered liquids consumed per cage normalized by the average body weight per cage. Two-way RM ANOVA post hoc Sídak’s Multiple Comparisons Test (b, d); T-test (c, e). ns = no significance, *p ≤ 0.05, ****p ≤ 0.0001. Values represent mean ± SEM.

Finally, in a separate set of mice, VNAA was compared to regular drinking water when mice were given one of the two drinking solutions ad libitum for a week (Figure 1Aiii). While there was a significant interaction between time and antibiotic solution (water n=9, VNAA n=12; 2-way RM ANOVA: interaction effect F[7,133]= 2.74, p=0.011), there was notably no main effect of antibiotic solution (F[1,19]=3.17, p=0.091). Šídák’s multiple comparison post hoc test found no significant differences in weight between water and VNAA groups at any time point (Figure 1D). Similarly, we found that cages receiving VNAA drank as much as cages receiving water when normalized to total body weight (Student’s unpaired t test: t[5]=1.04, p=0.593) (Figure 1E). Together, these data highlight VNAA as a viable antibiotic cocktail that is comparable to water in terms of body weight maintenance and volume consumed.

### 3.2 VNAA effectively depletes fecal microbes and does not induce anxiety-like behavior

We next tested whether the VNAA cocktail resulted in anxiety-like behavior using the open field test, as compared to mice receiving normal drinking water (Figure 2A). The total distance traveled in the open field was not significantly different between VNAA and water groups (2-way RM ANOVA: interaction effect F[1,28=0.415, p=0.600, main effect of time F[1,28]=5.87, p=0.013, and the main effect of solution F[1,19]=0.918, p=0.350) (Figure 2B). Likewise, there was no significant difference between groups for time spent in the center of the open field (2-way RM ANOVA: interaction effect F[2,38]=1.81, p=0.178, main effect of time F[2, 35]=2.12, p=0.139, and the main effect of solution F[1,19]=2.73, p=0.115) (Figure 2C). Similarly, behaviors were not significantly different after antibiotics compared to the pre-antibiotic baseline by Šídák’s multiple comparison post hoc test (Figures 2B-C).

**Figure 2.**
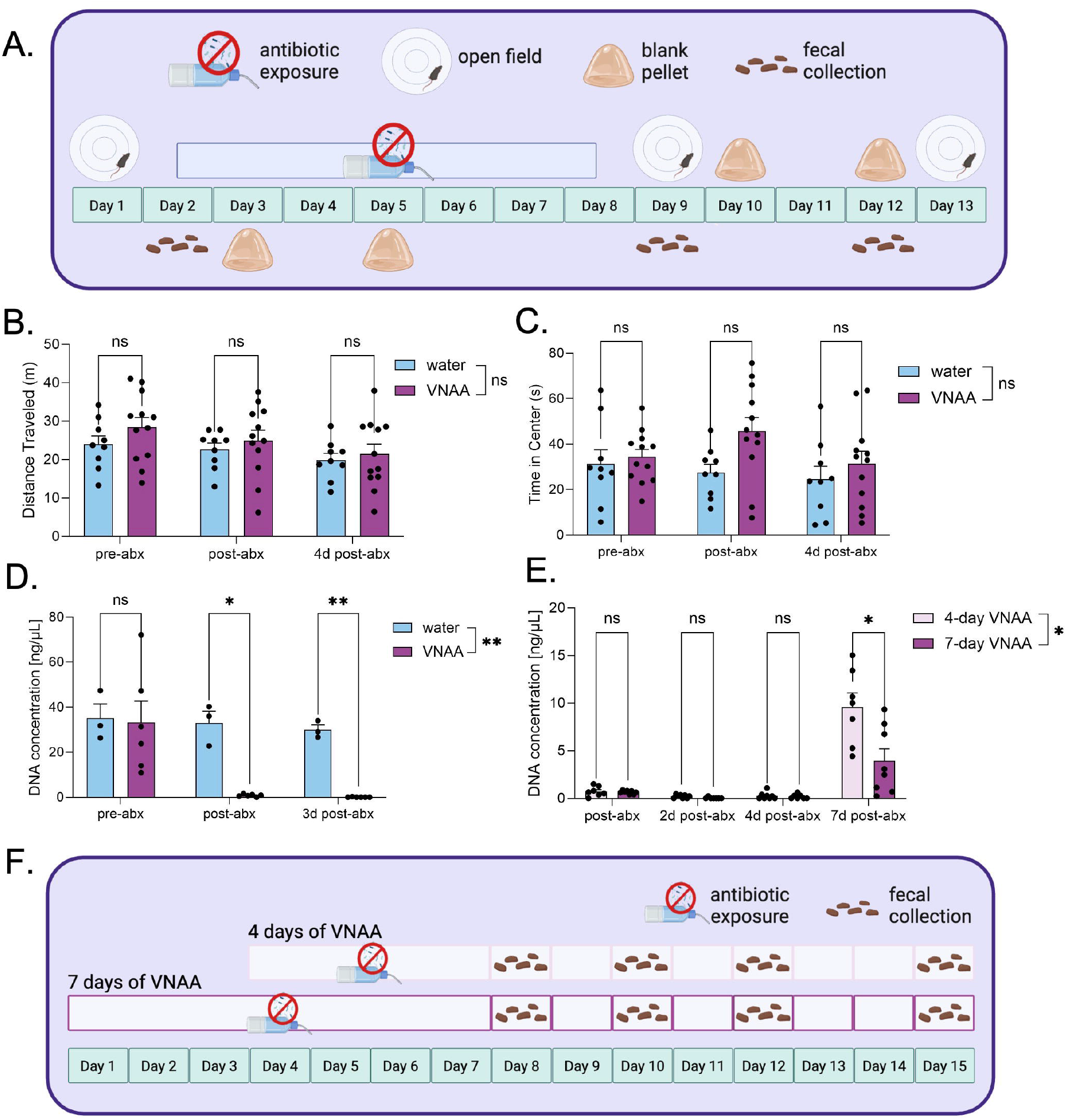
VNAA effectively depletes fecal DNA and does not elevate anxiety behaviors in mice. A: Experimental design for determining the antibiotic cocktail, VNAA’s ability to deplete gut microbial populations and measuring the behavioral stress response to VNAA. B-C: Quantifying overall distance and time spent in the center of an open field in mice before, after, and a few days after a week of VNAA. D. DNA concentration of fecal pellets in the open field experiment. E. DNA concentration of mice either drinking VNAA for four or seven days at different timepoints. F. Experimental design for determining an effective duration for VNAA administration. Two-way RM ANOVA post hoc Sídak’s Multiple Comparisons Test (b, d); Mixed-effects model (e). ns = no significance, **p ≤ 0.01, ****p ≤ 0.0001. Values represent mean ± SEM.

To assess the ability of VNAA to deplete the gut microbiome, we collected fecal samples from mice before, immediately following, and three days after a 7-day administration of VNAA or water control and compared microbial DNA concentration in the feces (Figure 2A). VNAA significantly depleted fecal microbial DNA concentration relative to the control (2-way RM ANOVA: interaction effect F[2,14]=3.94, p=0.044, main effect of timepoint F[1,8]=6.24, p=0.033, main effect of solution F[1,7]=17.5, p=0.004). Šídák’s multiple comparison post hoc indicated significantly less DNA in the VNAA-treated group (n=6) compared to the water-treated group (n=3) both on the final day of antibiotic treatment (p=0.026) and three days afterward (p=0.005) (Figure 2D).

We next compared the rate of microbiome recovery in mice receiving antibiotics for either four days (Tirelle *et al*., 2020) or one week (Figure 2F). The week-long regimen completely depleted fecal DNA concentrations two and four days after antibiotics (n=7; one-sample T test vs. hypothetical mean of 0: two days - t[6] = 1.00, p = 0.356, four days - t[6]=1.88, p=0.109). The four-day regimen reached depletion four days after antibiotics (n=7; one-sample T test vs. hypothetical mean of 0: two days - t[7]=3.19, p=0.015, four days - t[7]=1.90, p=0.099) (Figure 2E). Directly comparing the four and seven-day regimens, there was a significant interaction effect between time and treatment duration (REML: interaction effect F[3,38]= 8.64, p=0.0002, main effect of time F[1,13]= 46.2, p<0.0001, main effect of treatment duration F[1,14] = 8.05, p=0.013) (Figure 1E). Furthermore, one week following cessation of antibiotics, the group receiving antibiotics for 4 days showed significantly more fecal DNA, according to Šídák’s multiple comparison post hoc test (n=15, p=0.050) (Figure 2E). These data suggest that the four-day regimen is as effective at depleting fecal microbial populations as the seven-day regimen, but is less detrimental to microbiome recovery. We therefore used a four-day VNAA regimen for subsequent experiments in this study to better enable the engraftment and growth of the transplanted microbiomes.

### 3.3 Poopsicle preparation maintains anaerobic bacteria populations

Given the established increase in corticosterone following FMT via oral gavage (Gonzales *et al*., 2014), we next established an FMT procedure in which mice voluntarily consume fecal matter (“poopsicle”, herein referred to as pellet). This formulation contained a “fecal slurry” (see methods for details) that was frozen into aliquots. Pellet preparation occurred in an aerobic environment, potentially jeopardizing the survival of aerosensitive microbiota originating from the anaerobic environment of the colon. To determine whether our poopsicle method maintained anaerobic microbial stability (Figure 3A), we compared raw fecal matter collected from two individual donor mice to poopsicle pellets created from the same two individual donor mice. We plated swabs of the original starting fecal samples and our pellet homogenates, incubated in an anaerobic chamber with less than 25 parts per million O2 at 37°C for 72 hours, and collected bacterial colonies (Figure 3A). Finally, we sequenced the raw fecal matter (dark red bars), pellets (dark blue bars), plated raw fecal matter (light red bars), and plated pellets (light blue bars) to compare microbial content (Figures 3B-D). Anaerobic bacterial cultures demonstrated comparable DNA concentrations regardless of sample type (Figure 3B). Raw fecal and pellet samples also had similar DNA concentrations, and the DietGel contained minimal detectable bacterial DNA, suggesting that this was not a significant source of microbes (Figure 3B). The same trend was observed for the Shannon index, a measure of alpha diversity, assessed in samples from each donor mouse (Figure 3C). Comparing taxonomic parameters, phylum changes between raw vs. plated samples are larger than those measured between fecal vs. pellet samples (Figure 3D). In summary, plated fecal samples showed the most similar taxonomic composition when compared to plated pellet samples, while raw fecal and pellet samples showed the most similar taxonomic composition to each other.

**Figure 3.**
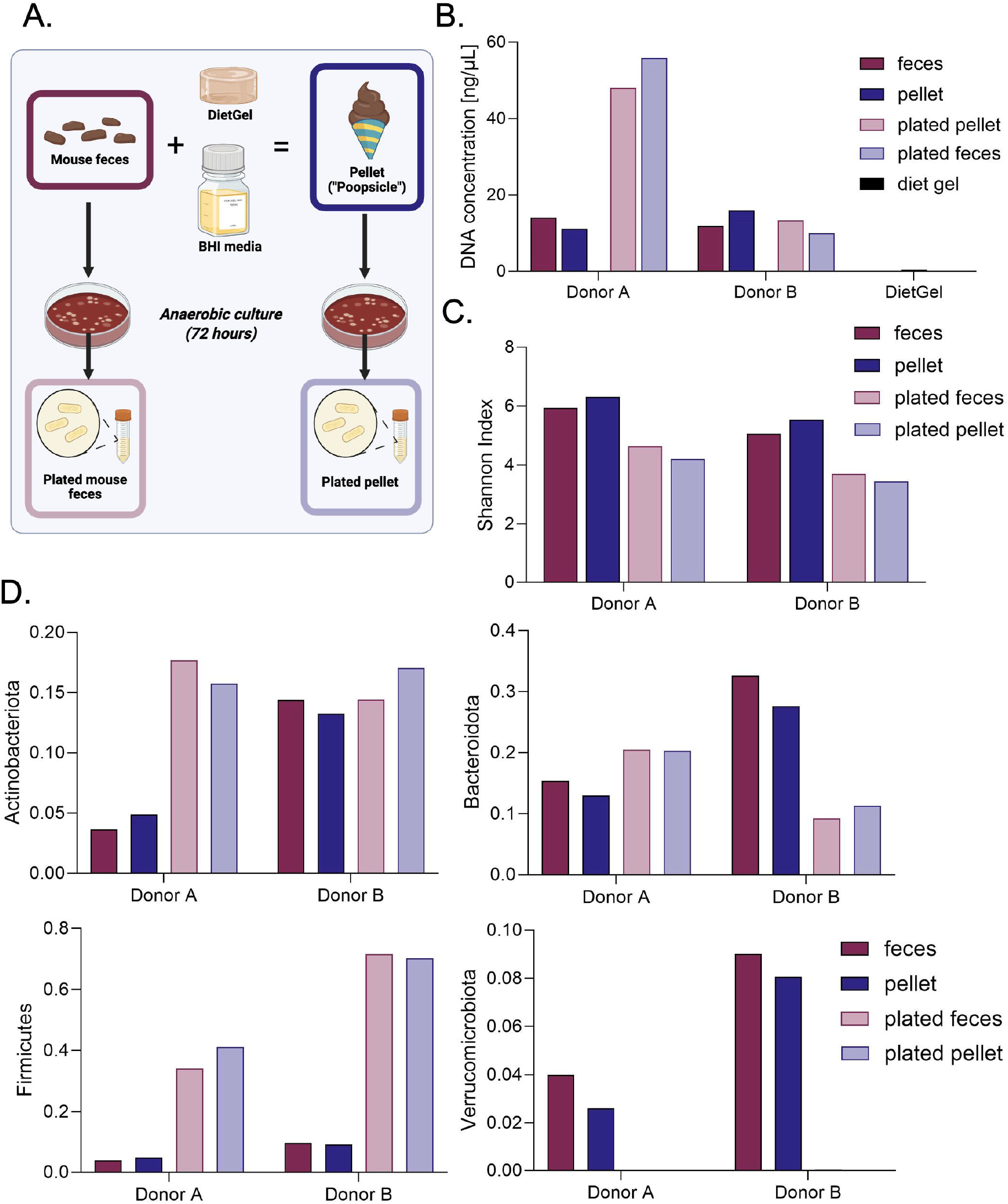
Anaerobic microbial communities are maintained during pellet preparation. A: Experimental design of pellet preparation and anaerobic culture. For each of the two donors we collected feces from, we measured the DNA concentration (b), Shannon index (c), and phyla (d) of their feces, the pellets made from their feces, and anaerobic cultures derived from each of the former sample types.

### 3.4 Delivery via poopsicle results in less circulating corticosterone than via oral gavage

Next, we investigated the recipient mouse stress response to consuming the pellets as compared to traditional oral gavage (Figure 4). We compared antibiotic-treated mice that received neither pellets nor oral gavage to mice receiving frozen pellets and to mice receiving oral gavage of fecal slurry (Figure 4A, see methods for additional experimental details). Twenty minutes following the third transplantation (either via pellet or oral gavage), blood was collected to measure plasma corticosterone. To account for additional or synergistic stress from antibiotics, we included an antibiotic control group, which received antibiotics but no transplantation (Figure 4A). The effect of FMT delivery method on circulating corticosterone was significant (one-way ANOVA: F[2,14]= 4.29, p=0.035). The concentration of corticosterone in mice receiving oral gavage (n=6) was significantly higher than the control group (n=5) (p=0.048, Tukey’s multiple comparisons post hoc test), while there was no significant difference between the control group and mice receiving the pellet (p=0.926). Additionally, the corticosterone levels of the mice receiving the pellet were marginally lower than those of the mice receiving gavage (p=0.078) (Figure 4B). Together, these results suggest that our novel voluntary pellet method is a less stressful alternative for delivering fecal microbiota transplants than oral gavage.

**Figure 4.**
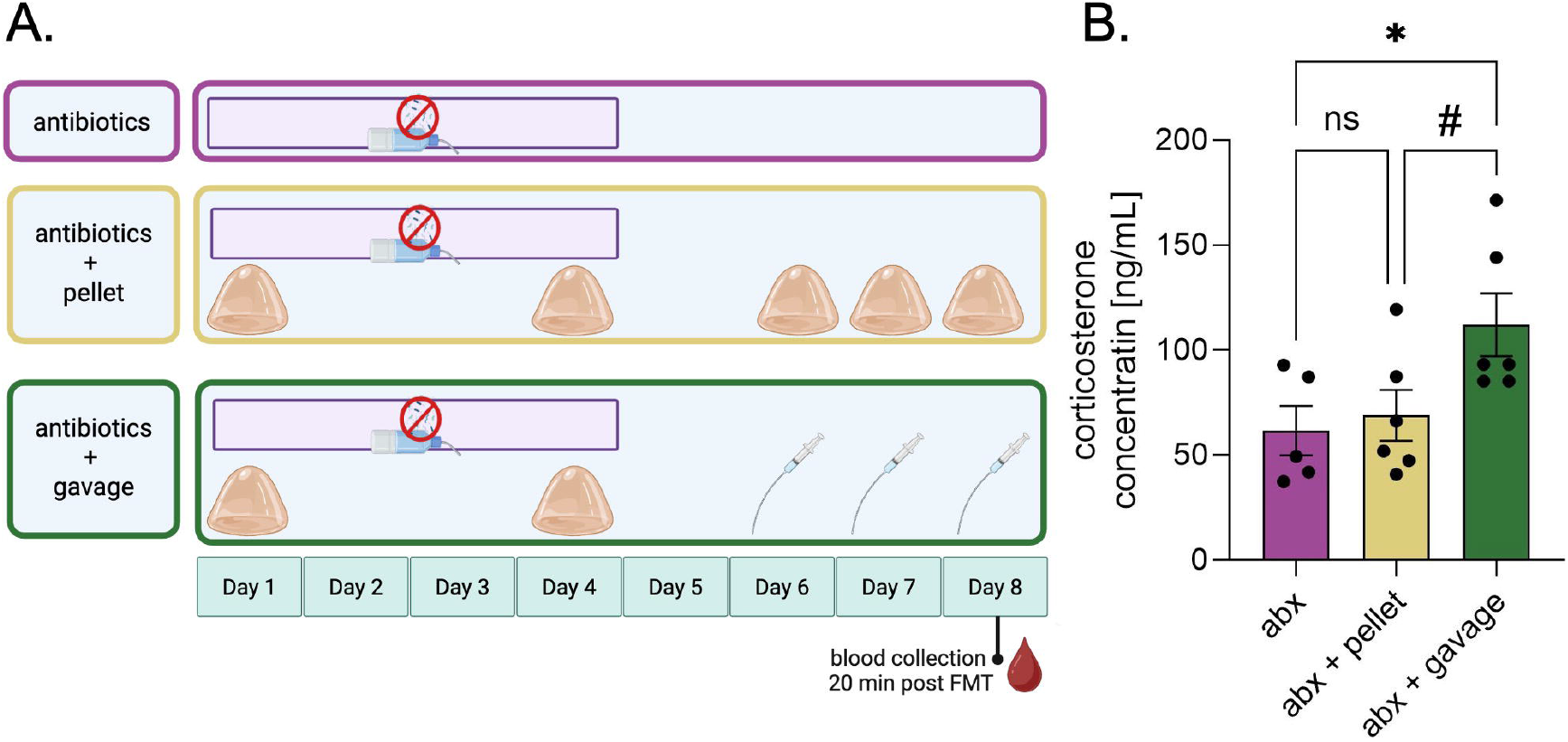
Delivery via pellet results in less circulating corticosterone than via oral gavage. A: Experimental design for determining the differences in stress between oral gavage and voluntary consumption as FMT delivery techniques. B: Concentration of circulating corticosterone in mice from each group. One-way ANOVA post hoc Tukey’s Multiple Comparisons Test (b); ns = no significance, # = trending, and *p ≤ 0.05. Values represent mean ± SEM.

### 3.5 Microbiomes of FMT recipients shift toward the donor

To evaluate the efficacy of the poopsicle pellet FMT method, we finally assessed the ability of our FMT method to shift the microbiome of recipient mice toward that of feces collected from donor mice fed a high-fat, high-sugar diet (see methods for additional experimental details) (Figure 5A). High-fat, high-sugar diets alter the gut microbiome, often promoting the proliferation of bacteria that damage the intestinal tract, creating a proinflammatory environment (Guo *et al*., 2021; He *et al*., 2024) Donor mice had access to the high-fat, high-sugar dietary supplement for six days before we collected and consequently pooled their feces and homogenized them with Diet Gel and a microbe-stabilizing media. Pellets were administered to all recipient mice except for those in the antibiotic control group (AB_control), which received blank pellets (no feces) instead. To determine the necessary dosage to see an effect, we compared two FMT dosing schedules: after a four-day regimen of antibiotics followed by a washout day, one group of mice (FMT1) received one FMT for three consecutive days followed by a booster FMT a week later, while the other group received the same treatment and followed by two extra weekly booster, for a total of three booster FMTs (FMT3). Before, throughout, and five weeks following the first three-day FMT, we collected feces from recipient mice for sequencing to determine how their microbiomes shifted throughout the study and how long the new microbiome lasted. (Figure 5A).

**Figure 5.**
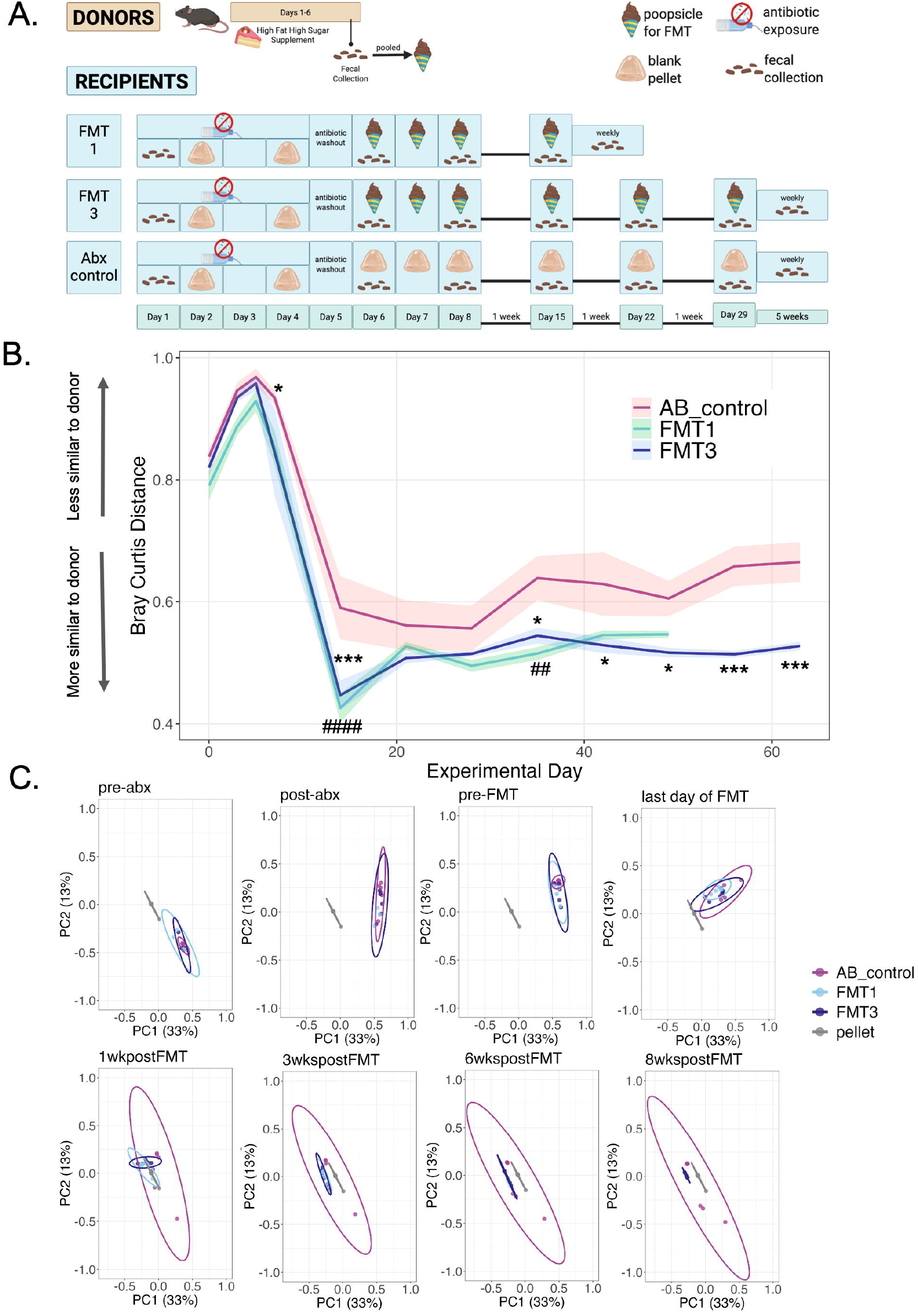
Microbiomes of FMT recipients shift toward the donor. A: Experimental design of donors and recipients. B: Univariate Bray-Curtis distance between each group and the average pellet across experimental days. C: PCoA plots mapping Bray-Curtis distances at various timepoints. Two-way RM ANOVA; comparisons between AB_control and FMT 1 are represented by # and comparisons between AB_control and FMT3 are represented by *; *p ≤ 0.05, ***p ≤ 0.001. Values represent mean ± SEM.

Following sequencing, we compared the average Bray Curtis distance of each collected fecal sample to each pellet sample at each timepoint (2-way RM ANOVA: interaction effect F[11,66]=1.50, p=0.152; main effect of timepoint F[11,66]=38.8, p<0.00001; main effect of treatment F[1,6]=11.3, p=0.015) (Figure 5B). Tukey’s multiple comparison post hoc test revealed that the distance to the pellet for the AB_control group was significantly different from the distance to the pellet for the FMT1 and/or FMT3 groups (Table 2), with FMT1 and FMT3 groups being closer to the original pellet material than the AB_control group (Figure 5B). Likewise, using principal coordinate analysis (PCoA) (Figure 5C) we found that all groups were significantly different from the pellet at each timepoint but that PERMDISP analysis demonstrated that the antibiotic control group consistently shows significant differences in dispersion (Table 3).

**Table 2:** Post-hoc analysis for Figure 5B: Tukey’s multiple comparisons.

| Groups Compared | Timepoint | Estimate | SE | Degree of Freedom | T ratio | P value | Annotation |
| --- | --- | --- | --- | --- | --- | --- | --- |
| AB_control - FMT1 | 1wkpostFMT | 0.164 | 0.0371 | 159 | 4.42 | **5.41E-05** | #### |
| AB_control - FMT3 | 1wkpostFMT | 0.143 | 0.0371 | 159 | 3.85 | **4.90E-04** | *** |
| FMT1 - FMT3 | 1wkpostFMT | -0.0210 | 0.0371 | 159 | -0.564 | 8.39E-01 |  |
| AB_control - FMT1 | 2wkspostFMT | 0.0341 | 0.0371 | 159 | 0.918 | 6.30E-01 |  |
| AB_control - FMT3 | 2wkspostFMT | 0.0534 | 0.0371 | 159 | 1.44 | 3.23E-01 |  |
| FMT1 - FMT3 | 2wkspostFMT | 0.0194 | 0.0371 | 159 | 0.521 | 8.61E-01 |  |
| AB_control - FMT1 | 3wkspostFMT | 0.0616 | 0.0371 | 159 | 1.66 | 2.24E-01 |  |
| AB_control - FMT3 | 3wkspostFMT | 0.0420 | 0.0371 | 159 | 1.13 | 4.97E-01 |  |
| FMT1 - FMT3 | 3wkspostFMT | -0.0196 | 0.0371 | 159 | -0.529 | 8.57E-01 |  |
| AB_control - FMT1 | 4wkspostFMT | 0.123 | 0.0371 | 159 | 3.32 | **3.14E-03** | ## |
| AB_control - FMT3 | 4wkspostFMT | 0.0942 | 0.0371 | 159 | 2.54 | **3.23E-02** | * |
| FMT1 - FMT3 | 4wkspostFMT | -0.0292 | 0.0371 | 159 | -0.786 | 7.12E-01 |  |
| AB_control - FMT1 | 5wkspostFMT | 0.08371 | 0.0371 | 159 | 2.254 | 6.54E-02 |  |
| AB_control - FMT3 | 5wkspostFMT | 0.100 | 0.0415 | 159 | 2.41 | **4.44E-02** | * |
| FMT1 - FMT3 | 5wkspostFMT | 0.0164 | 0.0415 | 159 | 0.397 | 9.17E-01 |  |
| AB_control - FMT1 | 6wkspostFMT | 0.0587 | 0.0371 | 159 | 1.58 | 2.57E-01 |  |
| AB_control - FMT3 | 6wkspostFMT | 0.0887 | 0.0371 | 159 | 2.39 | **4.73E-02** | * |
| FMT1 - FMT3 | 6wkspostFMT | 0.0300 | 0.0371 | 159 | 0.807 | 6.99E-01 |  |
| AB_control - FMT3 | 7wkspostFMT | 0.144 | 0.0371 | 159 | 3.88 | **1.53E-04** | *** |
| AB_control - FMT3 | 8wkspostFMT | 0.137 | 0.0371 | 159 | 3.70 | **2.97E-04** | *** |
| AB_control - FMT1 | last day of FMT | 0.0814 | 0.0371 | 159 | 2.19 | 7.55E-02 |  |
| AB_control - FMT3 | last day of FMT | 0.0892 | 0.0371 | 159 | 2.40 | **4.55E-02** | * |
| FMT1 - FMT3 | last day of FMT | 0.00782 | 0.0371 | 159 | 0.210 | 9.76E-01 |  |
| AB_control - FMT1 | post-abx | 0.0589 | 0.0455 | 159 | 1.29 | 4.00E-01 |  |
| AB_control - FMT3 | post-abx | 0.0116 | 0.0389 | 159 | 0.298 | 9.52E-01 |  |
| FMT1 - FMT3 | post-abx | -0.0473 | 0.0470 | 159 | -1.01 | 5.74E-01 |  |
| AB_control - FMT1 | pre-abx | 0.0474 | 0.0389 | 159 | 1.22 | 4.45E-01 |  |
| AB_control - FMT3 | pre-abx | 0.0178 | 0.0389 | 159 | 0.4583 | 8.91E-01 |  |
| FMT1 - FMT3 | pre-abx | -0.0295 | 0.0407 | 159 | -0.726 | 7.49E-01 |  |
| AB_control - FMT1 | pre-FMT | 0.0392 | 0.0407 | 159 | 0.964 | 6.01E-01 |  |
| AB_control - FMT3 | pre-FMT | 0.0103 | 0.0407 | 159 | 0.254 | 9.65E-01 |  |
| FMT1 - FMT3 | pre-FMT | -0.0289 | 0.0407 | 159 | -0.710 | 7.58E-01 |  |

**Table 3:** Analysis for Figure 5C: PERMANOVA Significance (q<0.05)

|  | AB_control vs Pellet | FMT1 vs Pellet | FMT3 vs Pellet |
| --- | --- | --- | --- |
| Pre-abx | **<0.001** | **0.001** | **0.001** |
| Post-abx | **<0.001** | **0.004** | **<0.001** |
| Pre-FMT | **<0.001** | **0.001** | **<0.001** |
| Last day of FMT | **<0.001** | **<0.001** | **<0.001** |
| 1wkpostFMT | **<0.001** | **<0.001** | **0.001** |
| 2wkspostFMT | **<0.001** | **<0.001** | **<0.001** |
| 3wkspostFMT | **<0.001** | **<0.001** | **<0.001** |
| 4wkspostFMT | **<0.001** | **<0.001** | **<0.001** |
| 5wkspostFMT | **<0.001** | **<0.001** | **0.001** |
| 6wkspostFMT | **<0.001** | **<0.001** | **<0.001** |
| 7wkspostFMT | **<0.001** | - | **<0.001** |
| 8wkspostFMT | **<0.001** | - | **<0.001** |

**Table 4:** Analysis for Figure 5C: PERMADISP Significance (q < 0.05)

|  | AB_control vs Pellet | FMT1 vs Pellet | FMT3 vs Pellet |
| --- | --- | --- | --- |
| Pre-abx | **<0.001** | **0.031** | **0.014** |
| Post-abx | **<0.001** | 0.065 | **<0.001** |
| Pre-FMT | **0.0017** | **0.003** | **<0.001** |
| Last day of FMT | **0.0015** | 0.670 | 0.095 |
| 1wkpostFMT | **0.004** | 0.136 | 0.162 |
| 2wkspostFMT | **<0.001** | 1.000 | 1.000 |
| 3wkspostFMT | **<0.001** | 0.589 | 1.000 |
| 4wkspostFMT | **0.004** | 0.522 | 1.000 |
| 5wkspostFMT | **<0.001** | 1.000 | 1.000 |
| 6wkspostFMT | **0.011** | 1.000 | 0.836 |
| 7wkspostFMT | **0.011** | - | 1.000 |
| 8wkspostFMT | **0.001** | - | 1.000 |

## 4 Discussion

The primary goal of this work was to develop and validate a low□stress fecal microbiota transplant (FMT) protocol that can be used to probe causal links in the microbiome□gut□brain axis. We first screened several antibiotic cocktails for tolerability and identified a vancomycin□neomycin□ampicillin□amphotericin□β mix that mice ingested voluntarily without weight loss. Unlike metronidazole, which has a bitter flavor and contributes to aversive drinking behavior, our regimen effectively depleted gut bacteria (including anaerobes) while preserving animal welfare. Next, we delivered the transplant as frozen pellets that mice voluntarily consumed. This approach eliminated the restraint and discomfort associated with oral gavage, a procedure known to raise corticosterone and confound stress□related readouts. In our acute stress assay, plasma corticosterone was markedly lower after pellet□based FMT than after gavage, indicating that the new method reduces the physiological stress response that can negatively affect behavioral outcomes. Moreover, the microbial composition of recipients remained closely aligned with the donor pellet signature for up to six weeks, as shown by Bray□Curtis analyses, demonstrating durable engraftment without the need for repeated invasive dosing. Thus, we have developed an accessible and low-cost fecal microbiota transplant protocol that minimizes treatment-induced stress.

One way to make FMT studies more accessible is to use antibiotics as the microbe-depletion method to clear the native gut flora instead of germ-free mice, which require a sterile and thus highly regulated facility. Metronidazole is a common antibiotic used to clear native gut microbiota in preparation for an FMT due to its cytotoxicity to anaerobes, the primary type of microbe in the oxygen-deficient gut (Löfmark, Edlund and Nord, 2010). However, we observed that animals have a strong aversion to metronidazole when administered ad libitum and lose a significant amount of body weight by refusing to drink (Figure 1B). Antibiotic cocktails with metronidazole are often combined with sucrose (Abt *et al*., 2012; Emal *et al*., 2017) and/or artificial flavoring such as Kool-Aid (Kang *et al*., 2008; Baldridge *et al*., 2015) to mask the antibiotic’s bitter taste and encourage drinking. However, this strategy can have confounding effects, especially in behavioral experiments (Richardson and McNally, 2003; Jones *et al*., 2005). Furthermore, artificial flavoring does not always resolve the refusal of animals to drink; thus, some researchers similarly remove metronidazole from their antibiotic regimen (Ogawa *et al*., 2020). Moreover, metronidazole can cross the blood-brain barrier and can result in encephalopathy, which can also confound neuropsychiatric outcomes depending on the disorder and brain region of interest, providing further justification for its removal from microbiota-depleting antibiotic cocktails (Agarwal *et al*., 2016).

As an alternative, we delivered a cocktail containing standard antibiotics, vancomycin, neomycin, and ampicillin. Vancomycin suppresses anaerobic gram-positive bacteria by inhibiting the synthesis of glycopeptide, a main component of bacterial cell walls (Koyama, Inokoshi and Tomoda, 2012; Tyrrell *et al*., 2012). Ampicillin and Neomycin have broad-spectrum activity against gram-positive and negative bacteria, including anaerobes, compensating for the lack of metronidazole (Jana and Deb, 2006; Snydman *et al*., 2018). Antibiotics for FMTs do not target fungal microbes residing in the gut (Kennedy, King and Baldridge, 2018); thus, for comprehensive treatment, we added the antifungal amphotericin-β, which targets key components of fungal cell membranes, leading to pore formation and cell death (Brajtburg *et al*., 1990). The combination of these antimicrobials thoroughly and broadly depletes microbial DNA in fecal samples, making it an effective method for drastically reducing the native microbiota without the need for metronidazole.

Antibiotic use is closely linked to several stress-related disorders through the gut-brain axis and can have various effects on behavioral outcomes (Barlow *et al*., 2022; Dinan and Dinan, 2022), which is why we measured the effect of antibiotics on anxiety-like behaviors in the Open Field test. Although seven days of VNAA did not significantly change anxiety-like or exploratory behaviors, we wanted to limit the effects of long-term antibiotic use on neuropsychiatric outcomes by limiting the duration of antibiotic exposure (Kennedy, King and Baldridge, 2018). In addition to impacting stress outcomes, prolonged antibiotic use could require more than one washout day to clear residual antibiotics and delay engraftment of the transplanted microbiome. We chose the four-day antibiotic regimen based on a study that demonstrated robust and consistent depletion of fecal microbial DNA after four days of twice-daily antibiotic administration (Tirelle *et al*., 2020). Tirelle et al. also noted that administering antibiotics for over seven days did not significantly change the amount of microbial DNA, indicating a floor effect, which was replicated in our data (Figure 2E). Therefore, four days of VNAA followed by one washout day were sufficient for microbiota clearance and limited confounding stress effects.

Considering stress outcomes also inspired the creation of an alternative to oral gavage for FMT delivery. Although we measured the acute stress response of our novel “poopsicle” technique as less stressful than that of oral gavage, oral gavage may result in limited side effects when performed by experienced technicians. One study found no differences in long-term cortisol levels between control and gavage animals; however, gavage animals had 15% mortality, with the mice having lesions and inflammation in the pleura, pericardium, and esophageal tissue (Arantes-Rodrigues *et al*., 2012). With animal models of stress or injury, additional treatment-induced acute or chronic inflammation can exacerbate the condition being studied, thus confounding results. Other studies have circumvented the need for oral gavage in FMT procedures; some facilitate the fecal transplantation between mice by simply cohousing separately treated animals or transferring cage bedding, which contains fecal droppings (Parker *et al*., 2022; Sun *et al*., 2023; Zhang *et al*., 2023; Urtecho *et al*., 2024). While this might be sufficient for other disease models, stress is transferred through olfactory avenues as well (Gómez-Sotres *et al*., 2024). Pheromones saturating the bedding can trigger the accessory olfactory bulb and the vomeronasal organ, both of which are implicated in conditioned fear responses (Carew *et al*., 2018). Cohousing introduces further complications with the added social effect of stress (Sterley *et al*., 2018; Horvath *et al*., 2025); other studies find that when a stressed mouse is cohoused with a non-stressed mouse, a mixed phenotype emerges within the cage, which is typically a median of the two ends of the spectrum (Steger *et al*., 2020). There are several studies that use sugar-coated gavages to incentivize the mice to receive the gavage (Hoggatt *et al*., 2010); however, this still risks damage to the upper gastrointestinal tract if the gavage needle is not carefully guided down the esophagus. Thus, while pellet-based FMT delivery may be an unorthodox choice, this choice was informed by careful consideration of confounding variables.

Similar to our choice to use antibiotics over germ-free mice, our FMT delivery procedure offers an accessible method easily implemented by rodent researchers, even those without access to anaerobic culturing infrastructure. The preparation of the pellet (poopsicle) maintained viable anaerobic populations that could be cultivated (Figure 3), likely due to minimal exposure to oxygen during pellet preparation, before freezing. Voluntary consumption of pellets shifted the microbiomes of antibiotic-treated recipient mice towards that of the donor microbiota (Figure 5D). The most similar microbiota between donor and FMT mice occurred one week after the first three FMT administrations (Day 15). The fact that the microbiome of the recipient FMT mice became more dissimilar from the pellet after this timepoint may indicate a need to deliver the first FMT booster sooner or a need for more boosters. Interestingly, the antibiotic control group also shifted toward the pellet after antibiotics. This observation could be attributed to antibiotics and a high-sugar, high-fat diet shifting microbiomes toward dysbiosis, resulting in altered neurotransmitter and glucose metabolism, as previously observed in other studies (Guo *et al*., 2021; He *et al*., 2024; Zhao *et al*., 2025). Thus, the FMT and antibiotics could result in similar shifts in microbial composition. However, the faster, more sustained, and robust shift toward the pellet in both FMT groups compared to the antibiotic control indicates that the voluntary consumption of the pellets did contribute to the change of composition. Ultimately, these results further demonstrate the accessibility of our approach by negating the need for anaerobic culturing infrastructure.

Given that this is a first attempt at a new method, we recognize that there are several limitations. For example, the poopsicles were created using pooled samples. Although we sequenced multiple poopsicles to verify their unique microbial signatures, we did not sequence individual donor fecal samples due to sample volume limitations, which restricted our experimental units, although we increased our observational units (Walter *et al*., 2020). Furthermore, all experiments were performed in male mice only. Future iterations and improvements of this design will investigate the feasibility of the method in female mice and perform sequencing of individual donor feces before pooling to leverage strain-resolved metagenomics that will enable precise identification of donor-specific taxa and allow quantitative tracking of their engraftment in recipients (Ianiro *et al*., 2022).

In sum, we have demonstrated that an acute antibiotic exposure paired with a voluntarily consumed, frozen fecal pellet formulation provides a robust, low-cost, low□stress avenue for fecal microbiota transplantation in mice, achieving durable shifts in microbial signatures toward donors while markedly attenuating the corticosterone surge associated with conventional oral gavage. This “poopsicle” method can also be applied to downstream probiotic experiments in rodents, eliminating the need for oral gavage administration at several points in the preclinical pipeline. By reducing the acute stressor of restraint□based delivery, this protocol preserves the integrity of stress□sensitive behavioral and neurobiological readouts, thereby strengthening causal inference in microbiome□gut□brain axis investigations. Future iterations and improvements of this design will continue to address the need for accessible strategies to rigorously study potential causal effects without confounding results to deepen our understanding of the association between stress and the gut microbiome. Overall, we hope this study provides a robust template for stress researchers to reference when carefully designing experiments that will thoughtfully disentangle the causal effects of the gut microbiome on gut-brain axis disorders.

## 6 Abbreviations

FMT: fecal microbiota transplant
ELISA: enzyme-linked immunosorbent assay
VNAM: vancomycin, neomycin, ampicillin, amphotericin-β, and metronidazole
VNAA: vancomycin, neomycin, ampicillin, and amphotericin-β
VNBP: vancomycin, neomycin, ampicillin, bacitracin, pimaricin

## 7 Disclaimer

The views expressed in this scientific presentation are those of the author(s) and do not reflect the official policy or position of the U.S. government or the Department of Veterans Affairs.

## 8 Declarations

### 8.1 Ethics Approval and Consent to Participate

All animal experiments were conducted in accordance with Association for Assessment and Accreditation of Laboratory Animal Care guidelines and were approved by the VA Puget Sound Institutional Animal Care and Use Committee.

### 8.2 Availability of Data and Materials

The data in this study are available from the corresponding author upon reasonable request.

### 8.5 Author Contributions

The work presented here was carried out in collaboration among all authors. MT, BE, SMG, AAW, and AS contributed to the conception and design of the study. MT, RDV, BE, DC, AW, RD, BS, and AS collected and analyzed data. MT and AS wrote the first draft of the manuscript. All authors contributed to manuscript revision, read, and approved the final manuscript.

## 8.6 Acknowledgements

We would like to thank Cindy Pekow, DVM, Leandra Mosca, DVM, MS, and Traci J. Weber for veterinary care. We also thank the University of Washington Microbial Interactions and Microbiome Center (mim_c) for fostering collaborations and providing 16S sequencing support. Timelines and experimental design summaries were created with Biorender.com. Lastly, the authors honor the mice, without which this research would not have been possible.

